# Pain sensitivity does not differ between obese and healthy weight individuals

**DOI:** 10.1101/2020.06.05.136598

**Authors:** Nichole M. Emerson, Hadas Nahman-Averbuch, Robert C. Coghill

## Abstract

There is emerging evidence suggesting a relationship between obesity and chronic pain. We investigated whether pain-free obese individuals display altered pain responses to acute noxious stimuli, thus raising the possibility of greater pain sensitivity and potential susceptibility for chronic pain development. Psychophysical and anthropometric data were collected from 39 individuals with an obese body mass index (BMI) classification (BMI≥30) and 40 age/sex-matched individuals of a healthy BMI (BMI<24.9). Since BMI may be an inaccurate index of obesity, additional anthropometric parameters of central adiposity, and percent body fat (BF%) were examined. Pain responses to supra-threshold noxious heat and cold stimuli were examined. Subjects provided pain intensity and unpleasantness ratings to noxious heat (49°C) applied at varying durations (5s, 12s, 30s) and locations (ventral forearm/lower leg). Cold pain ratings, thresholds, and tolerances were obtained following immersion of the hand in a cold-water bath (0-2°C). Between-group differences in pain responses, as well as relationships between pain responses and obesity parameters were examined. Importantly, confounds that may influence pain such as anxiety, depression, impulsivity, sleepiness, and quality of life were assessed. No between-group differences in pain sensitivity to noxious heat and cold stimuli were found. After controlling for sex, no relationships were found between BMI, central adiposity, or BF% and pain responses to noxious heat or cold stimuli. These results indicate that obesity, BF%, and central adiposity have little influence on pain sensitivity in obese individuals. Accordingly, it is unlikely that obesity alone increases susceptibility for chronic pain development via amplification of nociceptive processes.

## 1. Introduction

Obesity and chronic pain are two separate, yet intricately intertwined conditions that are currently major U.S. public health concerns. Prevalence of obesity is at epidemic levels with more than 35% of adults being classified as obese [42]. Similarly, about 30% of the population currently suffer from chronic pain conditions [1]. Both obesity and chronic pain decrease quality of life and are often comorbid with additional conditions such as depression, anxiety, and or poor sleep quality [2, 12, 16, 25, 26, 35, 56, 59, 70].

Previous studies have shown that chronic pain and obesity are related, such that obese individuals have an increased risk of developing chronic pain and individuals with chronic pain have an increased risk of being obese [28, 39, 46, 48, 58]. Chronic pain conditions relating to obesity are not limited to load-bearing conditions such as chronic low back pain, musculoskeletal pain, or knee osteoarthritis, although these are quite common [5, 14, 18, 19, 27, 28, 44, 47, 55, 57, 73]. Obesity has been linked to increased odds of chronic migraine, fibromyalgia, neck pain, abdominal pain, and chronic widespread pain [19, 73]. In addition, Stone and Broderick found that obese individuals were 68% more likely to experience pain than healthy weight individuals [65]. Recently, a correlation between BMI and prescription opioids was found, demonstrating that the risk of receiving prescription opioids increased progressively with BMI [64].

Although obesity and chronic pain are related, it is unclear whether obese individuals, that are otherwise healthy, are more sensitive to experimental pain and perhaps primed to develop a chronic pain condition. Studies have demonstrated conflicting results regarding whether sensitivity to experimental pain is altered in obese individuals [11, 23, 29, 32, 49, 53, 75]. These conflicting results across studies are likely due to differences in methodologies. Importantly, a critical limitation of the majority of these studies is the use of only BMI to assess obesity. Although BMI is a widely used and acceptable measure for obesity, it has major limitations. Most notably, BMI cannot differentiate between fat mass and lean muscle mass [40]. Thus, individuals with low body fat, but high muscle mass, could be identified as obese. On the other hand, individuals with a high body fat could have a healthy BMI (termed “normal weight obesity” or “skinny fat” [43]). In addition, the outcome measure of pain sensitivity has been most often defined using pain thresholds, and there is a need to assess other measurements such as supra-threshold pain ratings of pain intensity and unpleasantness [69]. Finally, co-morbidities that are common in obese individuals that may influence pain sensitivity such as diabetes, anxiety, depression, and quality/quantity of sleep are not often adequately controlled.

This study aimed to evaluate differences in pain sensitivity between healthy weight and obese individuals using several measures of obesity (BMI, central adiposity, and BF%), as well as multiple types, locations, and durations of supra-threshold nociceptive stimuli. Importantly, variability associated with obesity-related comorbidities was removed by controlling for confounding factors in the study design and data analyses.

## 2. Materials and Methods

### 2.1. Subjects

Study recruitment and assessment for eligibility occurred in 99 subjects, in which 20 were excluded for not meeting enrollment criteria. Exclusion criteria included BMIs between 25-29.9 kg/m^2^, a history of chronic pain conditions, chronic disease conditions, psychiatric disorders, neurological disorders, diabetes, and current medication use. Psychophysical and anthropometric data were collected from a total of 79 healthy volunteers (40 females, 39 males) ranging in age from 18-66 years with a mean age of 30 ± 9 years. The distributions of race/ethnicity included 58 Whites, 13 African Americans, 4 Asians, 2 Hispanics, 1 Indian, and 1 Multi-racial. Subjects were placed into age/sex-matched groups based on having a healthy BMI (n = 40, BMI = 18.5-24.9 kg/m^2^) or an obese BMI (n= 39, BMI ≥ 30 kg/m^2^). Subjects gave written informed consent stating they understood that they would experience painful thermal stimulation, that the experimental procedures were clearly explained, and that they could withdraw at any time without prejudice. Wake Forest University School of Medicine Institutional Review Board approved all study procedures.

### 2.2. Anthropometric Data Collection

Measurements of height, weight, waist circumference (WC), and skin-fold thickness were obtained. Subjects were weighed using an electronic scale. Heights were collected using a standard wall mounted stadiometer. Waist circumference was obtained by measuring the distance around the waist at the umbilicus. WC/height was used to calculate waist/height ratio (WHR). Skin-fold measurements were obtained on the direct skin of the left tricep, bicep, subscapular, and suprailiac regions in accordance with a standard protocol, using a Lange skin-fold caliper (Beta Technology, Watertown, WI) [10]. The sum of skin-fold measurements was used to calculate body fat percentage [10].

### 2.3. Self-Report Questionnaires

To identify potential confounding factors influencing pain perception and exhibiting relationships with obesity, subjects completed the Spielberger State-Trait Anxiety Inventory [61], Beck Depression Inventory-II [4], Epworth Sleepiness Scale [20], Barratt Impulsiveness Scale-11 [45], and the Short Form 36 Health Survey [71]. Questionnaires were completed prior to sensory testing.

### 2.4. Psychophysical Data Collection Overview

Subjects underwent a brief sensory training session prior to sensory testing. Sensory testing included stimulation with short and long duration heat stimuli and the Cold Pressor Test. Heat stimuli were applied using a 16 x 16-mm TSA II thermal stimulator (Medoc, Ramat Yishai, Israel), with 35°C serving as baseline. In order to prevent sensitization/habituation, the thermal probe was moved to a different location at the termination of each stimulus. The Cold Pressor Test was delivered via an ice water bath kept between 0-2°C. Subjects rated pain intensity and unpleasantness using a visual analog scale (VAS; Paresian Novelty, Chicago, IL) anchored at 0 (no pain and not at all unpleasant) and 10 (most intense pain imaginable or most unpleasant imaginable) [51, 52]. Subjects were instructed to only provide a rating for painful stimuli and thus provided pain intensity and unpleasantness ratings of zero if no pain was perceived.

#### 2.4.1. Sensory Training

In order to familiarize subjects with heat stimuli and use of the VAS, a sensory training session occurred prior to testing. During training, heat stimuli (35°C, 43°C-49°C) were applied to the left ventral forearm with rise and fall rates of 6°C/s, a plateau duration of 5s, and an inter-stimulus interval of 30s. Each stimulus temperature was delivered 4 times for a total of 32 stimuli. Subjects provided VAS pain intensity and unpleasantness ratings at the termination of each stimulus.

#### 2.4.2. Sensory Testing

The experience of pain can vary based on stimulus duration [21] due to both physical and neurophysiological factors. Brief heat stimuli will produce limited heating of deeper portions of the skin and will recruit populations of nociceptors with terminals in the superficial aspects of the skin [74]. In contrast, intermediate duration stimuli may also be perceived as more painful due to deeper diffusion of heat into the skin, and subsequent recruitment of deeper primary afferents [13]. In addition, long duration high intensity stimulation may result in increased pain intensity and unpleasantness, as well as changes in primary afferent and dorsal horn mechanisms that reflect temporal summation in addition to deeper diffusion of heat into the skin [6, 21, 31, 38]. In order to fully assess differences in pain sensitivity that may emerge only in long duration stimuli, 3 different stimulus durations were employed.

##### 2.4.2.1. Short Duration Heat Stimuli (5s)

Short duration (5s) heat stimuli (35°C, 43°C, 45°C, 47°C, 49°C) were applied to the right ventral forearm with rise and fall rates of 6°C/s, a plateau duration of 5s, and an inter-stimulus interval of 30s. Each stimulus was randomly delivered 3 times for a total of 15 stimuli. Subjects provided VAS pain intensity and unpleasantness ratings at the termination of each stimulus. The mean ratings to the three 49°C stimuli were used for analyses in order to enable comparisons with stimuli of other durations.

##### 2.4.2.2. Intermediate Duration Heat Stimuli (12s)

Ten intermediate duration (12s) heat stimuli (49°C) were applied to the left lower leg and interleaved with eleven 35°C stimuli in a 6.8-minute series with rise and fall rates of 6°C/s, plateau durations of 12s, and inter-stimulus intervals of 12s. Subjects provided a VAS pain intensity and unpleasantness rating at the termination of the series. One subject did not complete this stimulation due to the presence of scar tissue on the left calf stimulation site.

##### 2.4.2.3. Long Duration Heat Stimuli (30s)

Four long duration (30s) heat stimuli (49°C) were applied to the right lower leg and interleaved with five 35°C stimuli in a 4.8 minute series with rise and fall rates of 6°C/s, plateau durations of 30s, and inter-stimulus intervals of 30s. Subjects provided a VAS intensity and unpleasantness rating at the termination of the series.

##### 2.4.2.4. Cold Pressor Test

Noxious cold is processed differentially from noxious heat, both peripherally and centrally [8, 22, 34, 37]. Diverse processing of noxious cold yields differences in the subjective experience of pain, both in pain ratings and qualitative descriptors [34, 54]. Noxious cold is often delivered via the cold pressor test, which evokes higher ratings of pain unpleasantness compared to noxious heat pain in healthy individuals and may better represent mechanisms of chronic pain [54]. Accordingly, the cold pressor task was employed to examine sensitivity differences to cold pain between healthy BMI and obese BMI individuals. The cold pressor test was performed once per subject. Subjects submerged their left hands in an ice water bath (0-2°C) up to wrist level for 120s or until pain tolerance occurred. In order to promote circulation and prevent a boundary layer of warmth from forming around the hand, subjects were instructed to continuously open and close their hands for the duration of the test. In addition, subjects were instructed to notify the experimenter when they first felt pain and to remove their hand when they could no longer tolerate the pain. The duration of submersion until pain presented was recorded as threshold. The duration of submersion before the subject removed their hand was recorded as tolerance. VAS ratings of pain intensity and unpleasantness were taken every 30s and at termination of the test. In the results, only the pain intensity and unpleasantness ratings at the termination of the cold pressor test (tolerance) are presented due to the relatively short duration of pain tolerance.

### 2.5. Statistical Analyses

Statistical analyses were performed in JMP, version 11 (SAS Institute Inc., Cary, NC).

Differences in potential confounding variables were assessed between groups. Student’s t-tests were used to examine differences in anthropometric data and self-report questionnaire data between healthy BMI and obese BMI groups.

Pain ratings, BF%, and WC can differ based on sex [15, 33, 72]. In order to examine independent and joint effects of obesity and sex on pain sensitivity, BF%, and WHR (which is calculated as WC/height), two-factor ANOVA’s (group x sex) were employed. Regression analyses were used to further examine the relationships between pain intensity and unpleasantness ratings and BMI, WHR, and BF% in order to characterize potential effects of obesity that might not have been captured by dichotomous categorization based on BMI.

A total of 25 subjects did not reach tolerance during the cold pressor test. To control for ceiling effects, tolerance data for those subjects were not included in analyses.

For analyses of primary outcome variables, pain intensity and unpleasantness, results that initially met the significance threshold of p<0.05, were Bonferroni corrected for Type 1 error inflation due to multiple comparisons by reducing the significance threshold to (0.05/4), thus controlling for testing 4 stimulus paradigms. Both uncorrected and corrected p-values are reported.

## 3. Results

### 3.1. Anthropometric Measurements

Mean values for anthropometric data are displayed in Table 1. Subjects were placed into age and sex matched groups based on healthy BMI (BMI = 18.5-24.9 kg/m^2^) or obese BMI (BMI ≥ 30 kg/m^2^). Mean BMIs of the healthy BMI and obese groups were 22 ± 1.74 kg/m^2^ and 36 ± 5.23 kg/m^2^ (mean ± SD), respectively, and differed significantly (t(77) = 15.10, p< 0.0001). In addition, mean WC differed significantly (t(77) = 12, p < 0.0001) between healthy BMI (22 ± 6.7) and obese BMI (104.3 ± 14.3) groups. BF% differed significantly between groups (F(1,75) = 164, p<0.0001) and sexes (F(1,75) = 84, p<0.0001). The obese group displayed greater BF% than the healthy BMI group and females displayed greater BF% than males. No interaction between group and sex was found (F(1,75) = 0.30, p = 0.61). Similarly, a two-factor ANOVA revealed significant differences in mean WHR (F(1,75) = 148, p<0.0001) between groups, but not between sexes (F(1,75) = 1.1, p = 0.3). Additionally, no interaction between group and sex was found for WHR (F(1,75) = 0.06, p = 0.81).

**Table 1.**
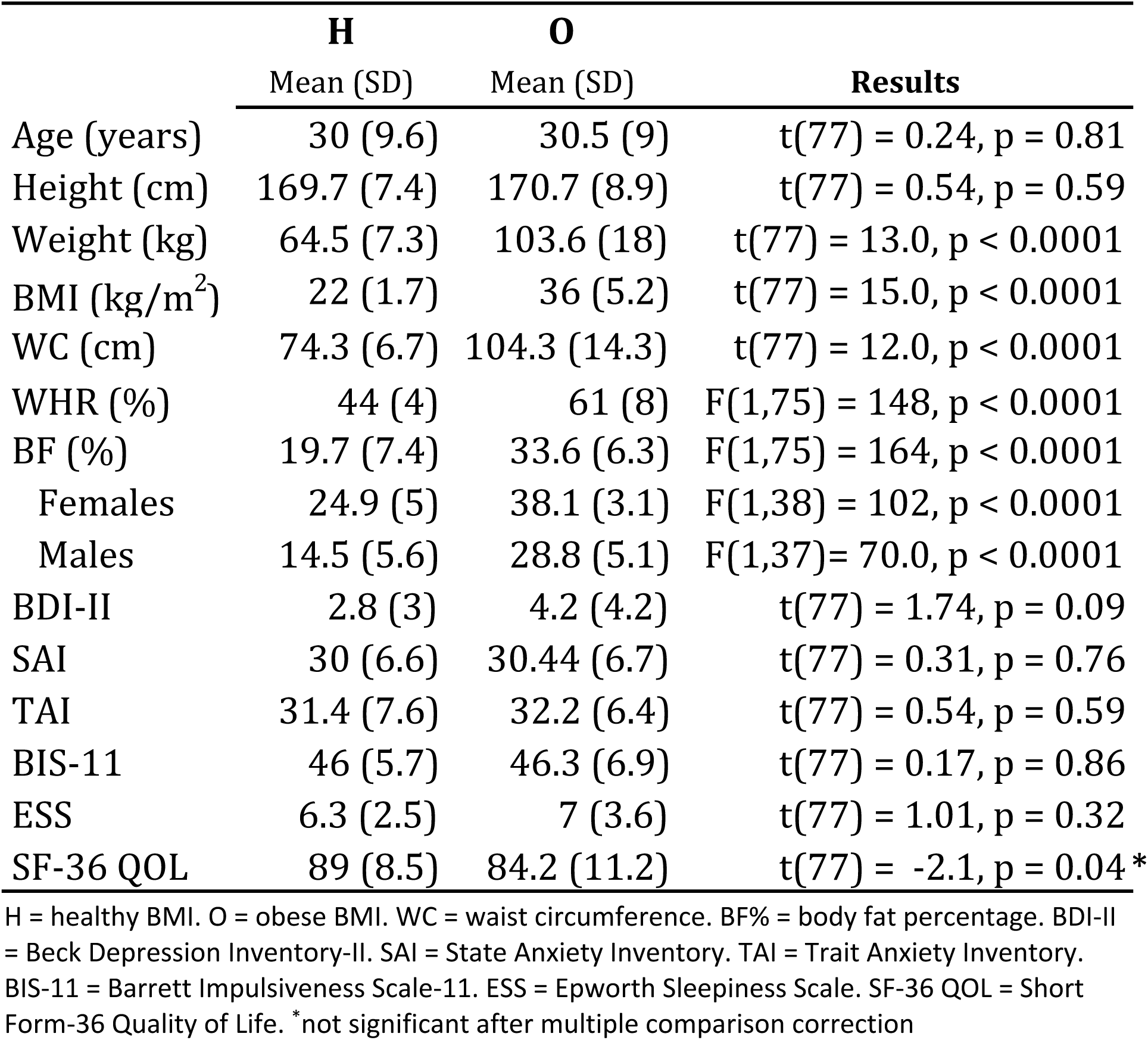
Demographic Data by Group

### 3.2. Self-Report Questionnaires

Factors such as anxiety, depression, impulsivity, sleepiness, and quality of life can co-vary with obesity and influence pain sensitivity. Accordingly, these factors were examined between groups to address possible confounds (Table 1). No significant between group differences were found for measures of state anxiety (t(77) = 0.31, p = 0.76), trait anxiety (t(77) = 0.54, p = 0.59), depression (t(77) = 1.74, p = 0.09), impulsivity (t(77) = 0.17, p = 0.86), or sleepiness (t(77) = 1.01, P = 0.32). Mean SF-36 quality of life scores did differ significantly (t(77) = -2.14, p = 0.04) between the healthy BMI (89 ± 9) and obese BMI (84 ± 11) groups.

### 3.3. Absences of Differences in Pain Sensitivity Between Healthy BMI and Obese BMI Groups

#### 3.3.1. Pain intensity

Perceived pain intensity can differ as a function of sex [33]. Accordingly, independent and joint contributions of obesity and sex were examined via 2-factor ANOVA’s. Between-group and between-sex comparisons are displayed in Figure 1/Table 2 and Table 3, respectively. Prior to Bonferroni correction, pain intensity ratings did not differ significantly between healthy BMI and obese BMI groups for short duration (5s) heat stimuli, intermediate duration (12s) heat stimuli, long duration (30s) heat stimuli, or noxious cold stimulus.

**Table 2.** Group differences in pain responses.

| Stimulus | Intensity<br>0-10 |  |  | Unpleasantness<br>0-10 |  |  |
| --- | --- | --- | --- | --- | --- | --- |
|  | H<br>Mean ± SD | O<br>Mean ± SD | F,p | H<br>Mean ± SD | O<br>Mean ± SD | F,p |
| Noxious Heat (5s) | 3.64 ± 2.31 | 3.85 ± 2.89 | F(1,75)=0.12, p=0.73 | 3.33 ± 2.31 | 3.88 ± 2.31 | F(1,75)=0.84, p=0.36 |
| Noxious Heat (12s) | 5.55 ± 2.46 | 5.42 ± 2.71 | F(1,74)=0.09, p=0.77 | 5.71 ± 2.54 | 5.69 ± 2.83 | F(1,74)=0.02, p=0.90 |
| Noxious Heat (30s) | 5.59 ± 2.72 | 5.64 ± 2.73 | F(1,75)=0.00, p=0.95 | 5.56 ± 2.90 | 5.78 ± 2.84 | F(1,75)=0.10, p=0.75 |
| Noxious Cold | 6.22 ± 2.59 | 6.42 ± 2.60 | F(1,75)=0.10, p=0.75 | 6.43 ± 2.79 | 6.96 ± 2.40 | F(1,75)=0.92, p=0.36 |
|  | Threshold |  |  | Tolerance |  |  |
|  | H<br>Mean ± SD | O<br>Mean ± SD | F,p | H<br>Mean ± SD | O<br>Mean ± SD | F,p |
| Noxious Cold | 15 ± 14 | 13 ± 10 | F(1,75)=0.61,<br>p=0.44 | 72 ± 42 | 69 ± 38 | F(1,75)=0.07, p=0.79 |

H = healthy BMI. O = obese BMI

**Table 3.**
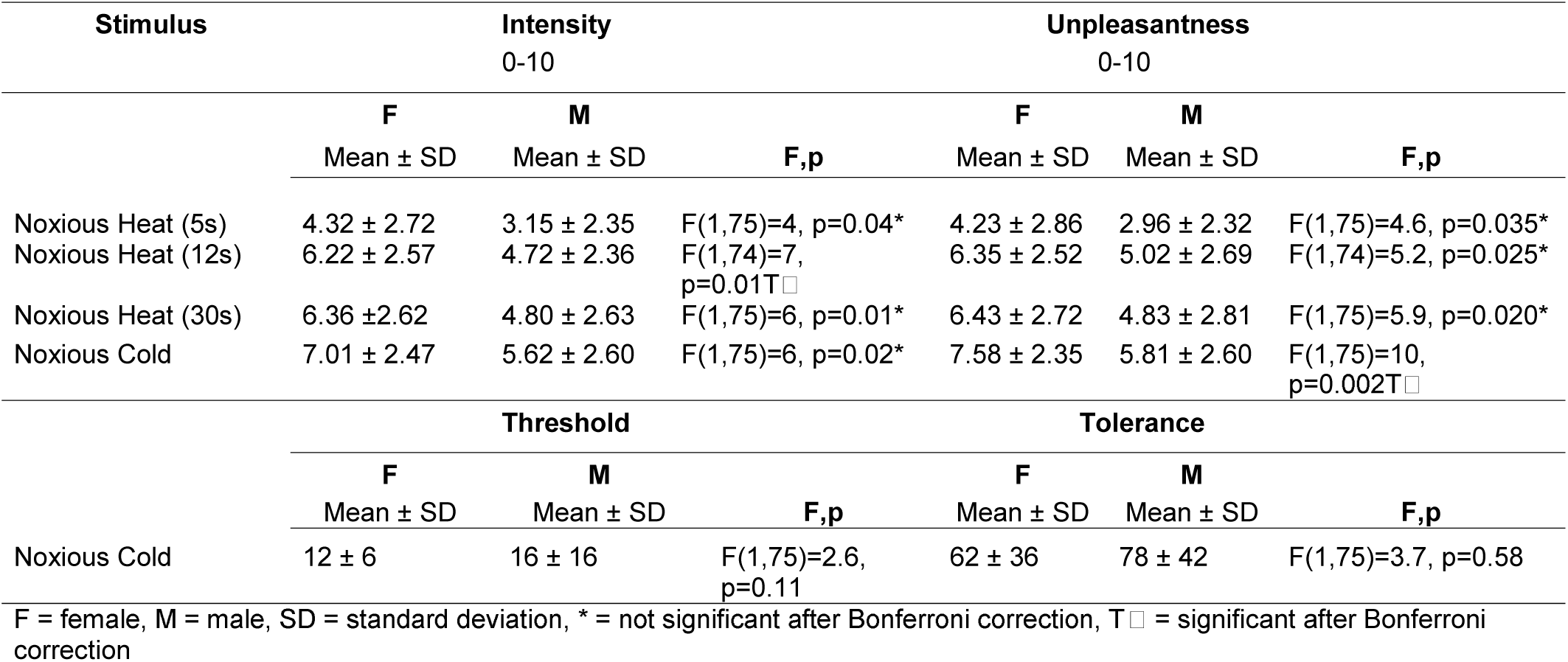
Sex differences in pain responses.

**Figure 1.**
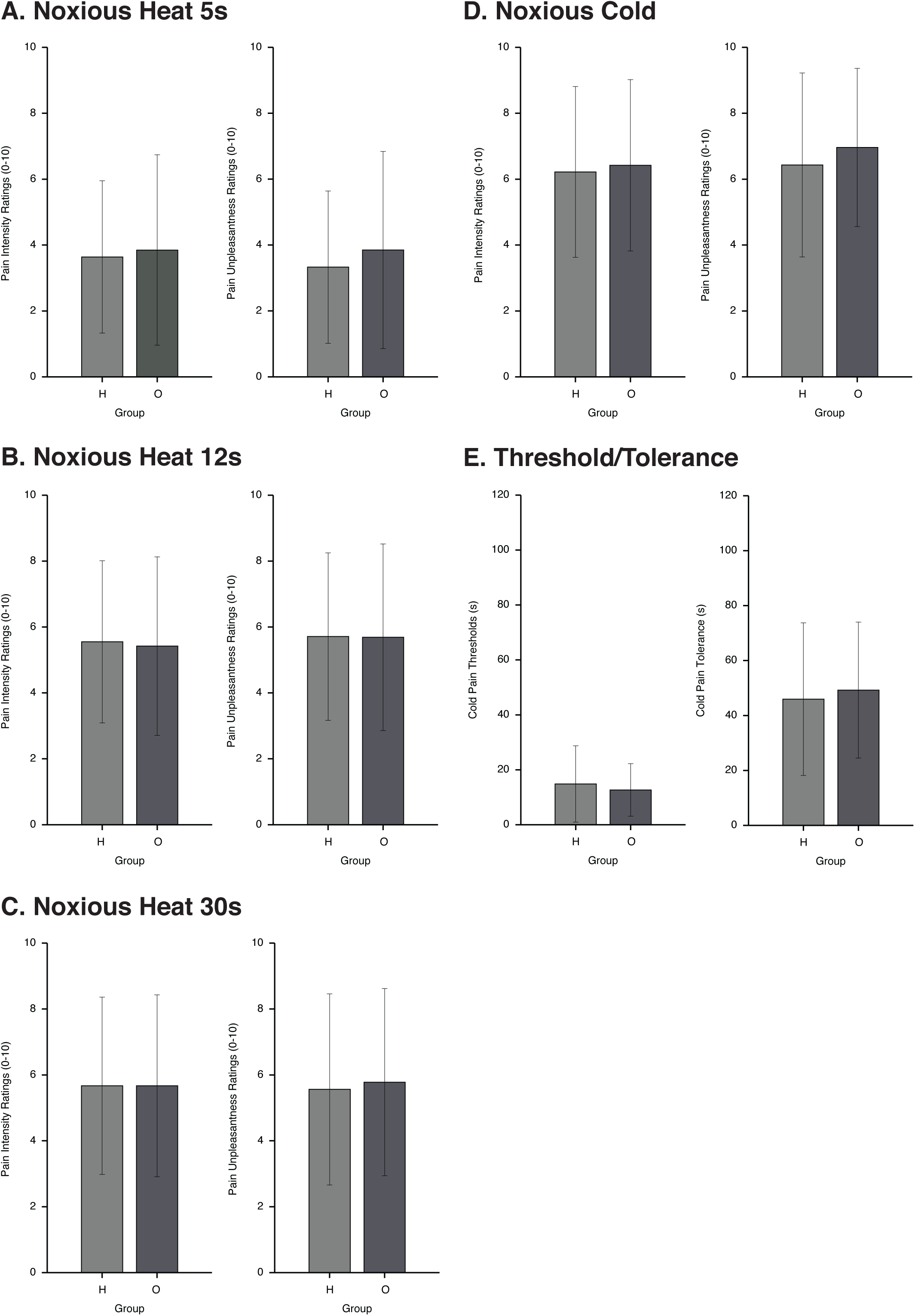
Pain evoked by experimental noxious stimuli does not differ between healthy weight and obese individuals. Investigation of suprathreshold heat (49°C) and cold stimuli (0-2°C) delivered to multiple locations at varying durations, found that pain responses did not differ significantly between healthy weight and obese weight groups. A-C. Between-group comparisons of pain intensity and unpleasantness ratings to noxious heat. D-E. Between group comparisons of pain intensity, pain unpleasantness, threshold, and tolerance to noxious cold delivered via the cold pressor test. A. Stimuli were applied for 5 seconds to the right ventral forearm. B. Stimuli were applied to the left calf for 12 seconds. C. Stimuli were applied to the right calf for 30 seconds. D. Cold pressor intensity and unpleasantness ratings were obtained at tolerance-driven termination of the test. E. Cold pressor threshold was recorded at the time the subject first reported pain. Cold pressor tolerance was recorded at the time the subject withdrew their hand from the test.

Conversely, prior to Bonferroni correction, sex differences in pain intensity were observed for short duration (5s) heat stimuli, intermediate duration (12s) heat stimuli, long duration (30s) heat stimuli, and cold stimulus (Table 3). Females displayed higher ratings of pain intensity than males. After correction for multiple comparisons, only pain intensity to intermediate duration (12s) heat stimuli remained significantly different between sexes. No interactions between group and sex were found for short duration (5s) heat stimuli, intermediate duration (12s) heat stimuli, long duration (30s) heat stimuli, or the cold stimulus.

#### 3.3.2. Pain unpleasantness

Prior to Bonferroni correction, pain unpleasantness ratings did not differ significantly between groups for short duration (5s) heat stimuli, intermediate duration (12s) heat stimuli, long duration (30s) heat stimuli, or cold stimulus.

As with pain intensity, prior to Bonferroni correction sex differences in pain unpleasantness were observed for short duration (5s) heat stimuli, intermediate duration (12s) heat stimuli, long duration (30s) heat stimuli, and the cold stimulus (Table 3). Females displayed higher pain unpleasantness ratings than males. After correcting for multiple comparisons, only pain unpleasantness for the cold stimulus remained significantly different between sexes. No interactions between group and sex were found for short duration (5s) heat stimuli, intermediate duration (12s) heat stimuli, long duration (30s) heat stimuli, or cold stimulus.

#### 3.3.3. Cold pain threshold and tolerance

No significant differences were found between groups for cold pain thresholds or tolerance (Table 2). In addition, no sex differences were observed in cold pain thresholds or tolerance (Table 3). Finally, no interaction was found between group and sex for cold pain thresholds or tolerance.

### 3.4. Relationships between Pain Sensitivity and BMI, WHR, and BF%

#### 3.4.1. BMI

Relationships between pain measures and BMI are shown in Table 4. After controlling for sex, no relationships were found between BMI and pain intensity ratings to short duration (5s) heat stimuli, intermediate duration (12s) heat stimuli, long duration (30s) heat stimuli, or the cold stimulus.

**Table 4.** Relationships between pain responses and obesity measures

| Stimulus | Intensity<br>0-10 |  |  |  | Unpleasantness<br>0-10 |  |  |  |
| --- | --- | --- | --- | --- | --- | --- | --- | --- |
|  | BMI | WHR | BF% |  | BMI | WHR | BF% |  |
| Noxious Heat (5s) | $R^2=.05$, $p=.91$ | $R^2=.05$, $p=.54$ | **F** $R^2=0$, $p=.92$ | **M** $R^2=0$, $p=.94$ | $R^2=.06$, $p=.74$ | $R^2=.06$, $p=.89$ | **F** $R^2=0$, $p=.80$ | **M** $R^2=0$, $p=.92$ |
| Noxious Heat (12s) | $R^2=.09$, $p=.67$ | $R^2=.10$, $p=.34$ | **F** $R^2=0$, $p=.55$ | **M** $R^2=.05$, $p=.20$ | $R^2=.07$, $p=.60$ | $R^2=.07$, $p=.35$ | **F** $R^2=.02$, $p=.38$ | **M** $R^2=.05$, $p=.19$ |
| Noxious Heat (30s) | $R^2=.08$, $p=.97$ | $R^2=.08$, $p=.99$ | **F** $R^2=.03$, $p=.30$ | **M** $R^2=0$, $p=.94$ | $R^2=.07$, $p=.81$ | $R^2=.07$, $p=.87$ | **F** $R^2=.04$, $p=.30$ | **M** $R^2=0$, $p=.94$ |
| Noxious Cold | $R^2=.07$, $p=.92$ | $R^2=.07$, $p=.86$ | **F** $R^2=0$, $p=.37$ | **M** $R^2=.02$, $p=.42$ | $R^2=.12$, $p=.81$ | $R^2=.12$, $p=.90$ | **F** $R^2=.04$, $p=.21$ | **M** $R^2=.01$, $p=.53$ |
|  | Threshold |  |  |  | Tolerance |  |  |  |
|  | BMI | WHR | BF% |  | BMI | WHR | BF% |  |
| Noxious Cold | $R^2=.05$, $p=.33$ | $R^2=.04$, $p=.45$ | $F(1,75)=2.6$, $p=0.11$ | | $62 \pm 36$ | $78 \pm 42$ | $F(1,75)=3.7$, $p=0.58$ | |

\*ratings recorded at termination of cold pressor test (0-120s). WHR = waist to height ration. F = female, M = male,

Likewise, no relationships were found between BMI and pain unpleasantness ratings to short duration (5s) heat stimuli, intermediate duration (12s) heat stimuli, long duration (30s) heat stimuli, or the cold stimulus.

Finally, BMI was not related to cold pain thresholds or tolerance.

#### 3.4.2. WHR

Relationships between pain measures and WHR are found in Table 4. After controlling for sex, no relationships were found between WHR and pain intensity ratings to short duration (5s) heat stimuli, intermediate duration (12s) heat stimuli, long duration (30s) heat stimuli, or the cold stimulus.

Similarly, no relationships were found between WHR and pain unpleasantness ratings to short duration (5s) heat stimuli, intermediate duration (12s) heat stimuli, long duration (30s) heat stimuli, or the cold stimulus.

Likewise, WHR was not related to cold pain thresholds or tolerance.

#### 3.4.3. Body fat percentage

Relationships between BF% and pain measures are shown in Table 4. After stratifying for sex, neither females nor males displayed a relationship between BF% and pain intensity ratings for short duration (5s) heat stimuli, intermediate duration (12s) heat stimuli, long duration (30s) heat stimuli, or the cold stimulus.

Similarly, no relationship was found in either sex between BF% and pain unpleasantness ratings for short duration (5s) heat stimuli, intermediate duration (12s) heat stimuli, long duration (30s) heat stimuli, or the cold stimulus.

Finally, no relationship was found in either sex between BF% and cold pain threshold or cold pain tolerance.

## 4. Discussion

Using noxious heat and cold stimuli across multiple body locations and durations of stimulation, we found no differences in supra-threshold pain intensity or unpleasantness ratings between obese and healthy BMI groups. Additionally, no between-group differences were found for cold pain thresholds or tolerance. Further, we found no relationships between obesity measures of BMI, WHR, or BF% and pain sensitivity. These results suggest that for otherwise healthy individuals, pain sensitivity does not differ based on obesity, or as a function of BMI, central adiposity or BF%. Accordingly, our findings indicate that obesity alone has little direct influence on pain sensitivity in healthy obese individuals.

Previous studies have produced conflicting results regarding altered pain sensitivity in obese individuals. Pradlier and colleagues found decreased nociceptive reflex thresholds (increased sensitivity) to electrical stimuli in obese individuals when compared to healthy weight individuals [49]. Similarly, a study examining pressure pain thresholds found increased sensitivity to mechanical stimuli in obese individuals when compared to healthy weight individuals [29]. Another study found lower pressure pain thresholds in the obese group compared to a healthy weigh group but no differences were found for heat and cold pain thresholds and tolerance [66]. Conversely, obese individuals displayed increased heat and cold pain thresholds (decreased sensitivity) on the fingers but not the toes [32]. Obese individuals also display increased pain thresholds and decreased pain ratings to noxious cold stimuli on the abdomen, but not on the forehead or hand [53]. Additionally, obese individuals exhibited increased pain thresholds to noxious heat stimuli on the abdomen, but no significant differences in pain ratings to a 1-minute 48°C stimulus [53]. Morbidly obese individuals also had higher electrical pain threshold and tolerance than healthy weight individuals but no differences in heat pain threshold and tolerance [68]. Contrary to the above studies that found some differences in pain sensitivity between obese and healthy weight individuals, other studies did not find such differences, which is in agreement with the results of the present study. In a large study (n=300), no relationships were found between BMI and pressure and heat pain thresholds as well as cold pain tolerance using the cold pressor test [36]. This study also included a thorough examination of covariates such as anxiety, depression, and quality of life. Our findings are also in agreement with another study that found no difference in electrical pain ratings between obese and healthy weight groups [23]. Although psychological data were not collected for that study, subjects were healthy and without underlying medical conditions or diabetes. Another recent study found no differences in pressure pain thresholds and the conditioned pain modulation response between healthy subjects that have normal BMI and high BMI [11]. In addition, no correlations were found between BMI and pressure pain thresholds [11]. Possible sources for the discordance across studies are differences in psychophysical and anthropometric measures, healthiness of obese subjects, and control of psychological factors that may influence pain.

There are several important features of the present study. First, the present study utilized a comprehensive exploration of anthropometric measures including BMI, central adiposity, and BF%. The majority of the previous studies relied solely on “% above ideal weight” or BMI as the marker of obesity. These measures have known disadvantage and may misclassify individuals that are athletic or with high muscle mass as obese or vice versa, misclassify individuals with high fat mass as normal weight [40, 43]. Thus, it is important to include other measures of obesity such as central adiposity, and BF%, as was done in the present study. In addition, the present study controlled for comorbidities and psychological factors that may influence pain sensitivity. Subjects in the present study were all healthy, without medical conditions, and not taking any medications. Previous findings of altered pain sensitivity in the obese may have been driven by the effects of confounding conditions on pain sensitivity, rather than the obesity itself [29, 32, 49]. In addition, previous studies have not accounted for possible psychological confounds relating to obesity that may influence pain, such as depression, anxiety, impulsivity, quality of life, and sleepiness [2, 35, 56, 59, 70]. In contrast, the present study addressed comorbidities and psychological factors in both the study design (exclusion criteria) and analysis (sub-clinical presentations). Of note, psychological factors that may contribute to shaping pain unpleasantness, such as anxiety and depression, also did not differ significantly between groups. In addition, the present study used supra-threshold stimuli to assess differences in pain sensitivity (as opposed to thresholds) and included ratings of pain unpleasantness. Pain thresholds and supra-threshold pain responses are substantially different measures and are poorly correlated within individuals [62]. Pain thresholds, by definition, measure the border between no pain/pain and are vulnerable to influence by subjective factors [63]. Moreover, while previous studies provided little information about levels or dimensions (intensity/unpleasantness) of the pain experience [63], the present study assessed both pain intensity and unpleasantness ratings. Pain unpleasantness represents the affective component of pain and is distinguished from the sensory component of pain [30, 50]. Since pain is a multi-dimensional experience consisting of sensory, affective, and cognitive components [30], it is important to assess pain unpleasantness in addition to pain intensity. To our knowledge, the present study is the first to examine whether pain unpleasantness is altered in obese individuals and whether it is related to BMI, central adiposity, or body fat percentage. No significant differences in pain unpleasantness were found between healthy weight and obese individuals and no relationships were found between pain unpleasantness and BMI, central adiposity or body fat percentage.

Importantly, our findings are consistent with what is known about peripheral nociceptive processes. Afferent fibers terminate in the epidermis, superficial to the subcutaneous hypodermis where adipose tissue is stored [60, 67, 76]. Therefore, it is unlikely that adipose tissue directly interferes with nociception via a blocking or buffering of nociceptor activation. Thus, no differences in pain sensitivity between healthy weight and obese individuals were found even for the short noxious heat stimuli. Interestingly, Price and colleagues have proposed that a decrease in afferent fiber density, due to skin stretching, may account for pain sensitivity differences in obese subjects [53]. A between-groups analysis found that obese subjects displayed differences in pain thresholds to stimuli delivered to the abdomen, but not the hand or forehead. The present study cannot be directly compared with those findings, as pain responses to abdominal stimulation were not measured. However, the finding of only abdominal alterations is consistent with the present study, suggesting that pain sensitivity is not systemically altered in obese groups.

Additionally, and in support of the present findings, it has been shown that pain sensitivity does not change after substantial weight loss following bariatric surgery [7]. Furthermore, a large investigation of pain measures in chronic back pain patients found no difference in pain severity or frequency between healthy weight, overweight, and obese groups [24]. This collection of evidence, coupled with the results from the present study, indicates that increased adiposity alone does not alter pain sensitivity in healthy individuals.

The mechanisms that link obesity and pain can be mechanical (excess weight on joints that can lead to injury and damage to the joints and thus, pain), behavioral (lower physical activity and higher rates of sleep disturbance in obese individuals) and physiological (secretion of pro-inflammatory cytokines) [9] Sex was also suggested to be a factor since it has a role in both pain sensitivity and obesity [3, 15, 17, 41]. In the present study sex differences were found for BF% and pain intensity ratings of heat stimulus (12 seconds) and cold stimulus. However, no effect of sex was found for the relationships between anthropometric measurements and pain sensitivity.

## 5. Limitations

Because this study was limited to healthy obese subjects, we did not examine whether pain sensitivity is altered in metabolically unhealthy obese individuals. However, these results indicate that if pain sensitivity is altered in metabolically unhealthy obese, it is most likely a by-product of a co-varying underlying condition and not the direct result of increased adiposity. Future studies should aim to delineate differences between obese individuals with and without chronic pain in an effort to uncover factors that may contribute to the increased risk of chronic pain development in obese individuals. In addition, we did not examine mechanical or electrical pain, and therefore cannot comment on whether those modalities display alterations in pain sensitivity.

## 6. Summary

Risk of chronic pain development is increased in obese individuals. Using multiple psychophysical and anthropometric measures, we investigated whether healthy obese individuals display alterations in pain sensitivity that may increase susceptibility to chronic pain development. We found that pain sensitivity was not altered in obese individuals regardless of testing modality, anatomical location of testing, or duration of stimulation. Additionally, we found no relationships between pain measures and BMI, central adiposity, or BF%. These results suggest that although chronic pain and obesity are related, increased adiposity does not in-and-of-itself alter nociception. It is therefore unlikely that obesity directly contributes to chronic pain development via an adiposity-related amplification of nociceptive processes.

## Acknowledgment

This study was funded by grant number R01NS039426 and R01NS085391.

